# Rigidifying a *de novo* enzyme increases activity and induces a negative activation heat capacity

**DOI:** 10.1101/2021.04.16.439788

**Authors:** SA Hindson, HA Bunzel, B Frank, DA Svistunenko, C Williams, MW van der Kamp, AJ Mulholland, CR Pudney, JLR Anderson

## Abstract

Conformational sampling profoundly impacts the overall activity and temperature dependence of enzymes. Peroxidases have emerged as versatile platforms for high value biocatalysis owing to their broad palette of potential biotransformations. Here, we explore the role of conformational sampling in mediating a *de novo* peroxidase’s activity. We demonstrate that 2,2,2-triflouoroethanol (TFE) affects the equilibrium of enzyme conformational states, tending towards a more globally rigid structure. This is correlated with increases both stability and activity. Notably, these effects are concomitant with the emergence of curvature in the temperature-activity profile, trading off activity gains at ambient temperature with losses at high temperatures. We apply macromolecular rate theory (MMRT) to understand enzyme temperature dependence data. These data point to an increase in protein rigidity associated with a difference in the distribution of protein dynamics between the ground and transition state. We compare the thermodynamics of the *de novo* enzyme activity to those of a natural peroxidase, horseradish peroxidase. We find that the native enzyme resembles the rigidified *de novo* enzyme in terms of the thermodynamics of enzyme catalysis and the putative distribution of protein dynamics between the ground and transition state. The addition of TFE apparently causes C45 to behave more like the natural enzyme. Our data suggest robust, generic strategies for improving biocatalytic activity by manipulating protein rigidity; for functional *de novo* protein catalysts in particular, this can provide more enzyme-like catalysts without further rational engineering, computational redesign or directed evolution.

---

Biocatalysis is central to the development of a sustainable chemical industry.^1^ Many natural enzymes are extremely proficient, highly specific, and can provide access to novel and efficient synthetic routes. Nonetheless, their biocatalytic applications are limited by the exquisite selectivity that enzymes display for their natural transformations. Creation of novel enzymes through *de novo* design or directed evolution provides a means to fill such gaps in the biocatalytic toolbox.^5^These engineering efforts often have significant effects on the scaffold dynamics,^6^ and a complete understanding of the molecular and dynamic changes wrought by engineering – particularly the effects on the thermodynamic drivers of catalysis – is required to rationally create better enzymes in the future.

Heme peroxidases are versatile biocatalysts that exploit their heme cofactor to form highly oxidative oxy-ferryl intermediates (Figure 1B).^6^ We have recently created the *de novo* heme peroxidase C45 (Figure 1A), a rationally designed four-helical bundle protein with high peroxidase activity.^7^ Notably, C45 activity approaches that of natural benchmarks such as horseradish peroxidase (HRP) but is limited by the lack of a catalytic proton-shuttling residue, which likely slows down both the initial H_2_O_2_ deprotonation and the subsequent cleavage of the peroxide O-O bond; this results in an elevated *K*_M_ for H_2_O_2_.^7^ The relative simplicity of these *de novo* designed helical bundles allows facile tailoring for specific purposes, e.g. by introducing site-specific functional moieties.^8–11^ Furthermore, they are often highly flexible, facilitating access to a broad equilibrium of conformational states for engineering.^12^ In contrast to many other heme-dependent enzymes, C45 catalyses a broad range of transformations such as dehalogenations and carbene-transfer reactions.^13^

**Figure 1.**
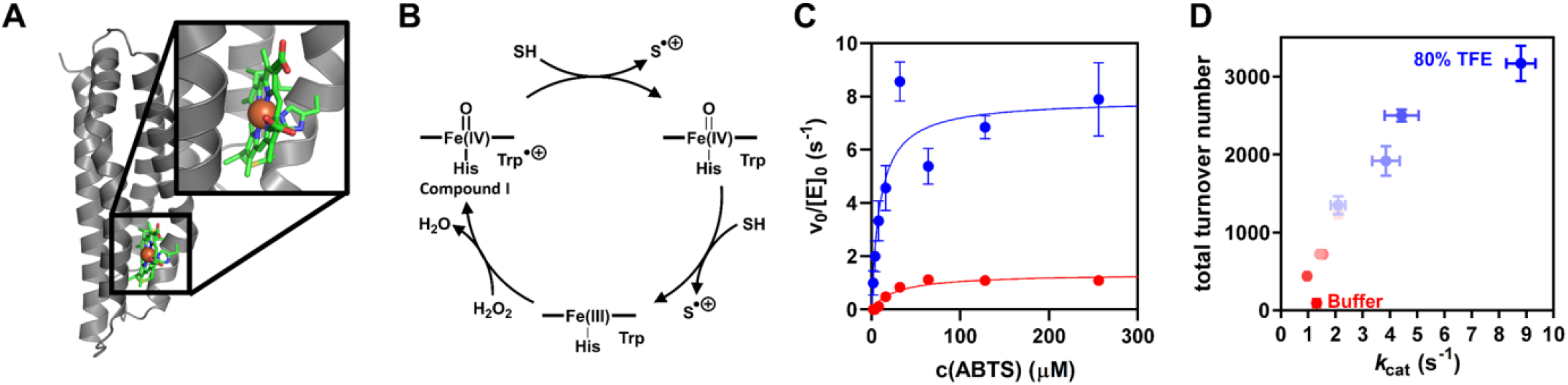
TFE boosts activity of *de novo* heme-peroxidase C45. **A**, The heme cofactor of C45 (green sticks, iron: brown sphere) gives rise to catalytic peroxidase activity **(B). C**, Addition of 80% TFE (blue) boosts oxidation of ABTS by C45 compared to buffer (red). **D**, TFE additionally increases the total turnover number (TTN) of C45 (red to blue: 0% to 80% TFE). *Conditions*, 25-200 nM C45, 100 µM H_2_O_2_, 100 mM KCl, 20 mM CHES, pH 8.6, 25 °C.

Understanding the relationship between protein dynamics and enzyme turnover holds great promise for the ability to engineer enzyme activity. Moreover, there are important fundamental questions regarding the relationship between the global protein flexibility and the conformational dynamics that are relevant to enzyme turnover. Recent efforts to understand the relationship between protein flexibility (the equilibrium of conformational states; dynamics) and enzyme turnover have employed macromolecular rate theory (MMRT)^12–14^ to infer differences in the distribution of vibrational modes between the ground and transition state. Evidence from a range of experimental and computational studies point to the involvement of networks of protein dynamics that extend throughout the protein and are not just localised to the immediate active site.^15–17^ MMRT quantifies the difference in distribution of vibrational modes between the ground and transition state ensemble as the heat capacity of catalysis (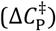) given by,

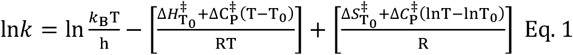

Where, 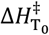 and 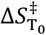 are the activation enthalpy and entropy at an arbitrary reference temperature *T*_0._ Where such curvature is present, values of 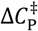 are typically negative for natural enzymes, manifesting as concave curvature in plots of ln*k versus T*.

Here we provide evidence that global rigidification of C45 increases enzyme activity, and also induces a difference in rigidity (dynamics) between the ground and transition state ensembles. By comparison to a natural enzyme, our data point to a complex interplay between enzyme activity, stability and protein dynamics. We discuss the implications for protein engineering principles that might leverage this emerging relationship. Our results point to a practical route of modulation of *de novo* enzyme dynamics, activity and thermal optima.

## Results and Discussion

### Trifluoroethanol increases peroxidase activity

When studying the effects of various organic solvents on C45 peroxidase activity, we discovered a ∼6-fold increase in activity for the oxidation of ABTS (2,2′-azino-bis(3-ethylbenzothiazoline-6-sulfonic acid) in 80% 2,2,2-trifluoroethanol (TFE) (Figure 1C) under limiting peroxide concentrations with *k*_cat_ = 1.3 ± 0.2 s^−1^ and 7.9 ± 0.9 s^−1^, respectively. We observe a very large increase in the total turnover number (TTN), with a ∼35-fold increase in 80% TFE compared to the absence of TFE (Figure 1D).

We have monitored the stability of the catalytic competency of C45 from steady-state progress curves as shown in Figure S1. At low temperatures, our progress curves are linear for > ten minutes, however at elevated temperatures there is apparent curvature. In the absence of trivial effects such as substrate depletion, this finding is typically indicative of enzyme inactivation/unfolding. From Figure S1, the curvature is rather more pronounced in the absence of TFE on the same timescale, suggesting that TFE stabilises the catalytically competent conformational state of C45. Our data therefore suggests that the reason for the higher TTN in the presence of TFE arises not just because of a faster rate of turnover and more stable intermediate, but also at least in part, a more stable enzyme. See below for further discussion on C45 stability with TFE.

Peroxidases form highly reactive oxyferryl intermediates that may oxidize and damage the protein and/or the heme cofactor,^6^ and the apparent increase in TTN suggests that in 80% TFE, C45 favours productive turnover over such detrimental off-pathway reactions. There is range of potential mechanisms through which we can envisage TFE acting, including; stabilisation of Compound I and/or prevention of heme degradation *via* protein rigidification, which inhibits the reaction with neighbouring side chains; ABTS binding more readily to the protein surface in TFE, similarly preventing a Compound I self-reaction.

To probe the mechanistic underpinning for the enhanced activity, we determined the kinetics of Compound I formation using pre-steady state kinetics (Figure 2A-D). H_2_O_2_ turnover in the presence of C45 proceeded with multiphasic kinetics, with the first phase relating to formation of an intermediate, and subsequent phases to its decay and heme degradation as indicated by an overall absorbance bleach. EPR spectroscopy allowed to electronically characterise that intermediate after freeze-quenching variable time following addition of H_2_O_2_. An anisotropic EPR signal at g = 2.0044 was observed, its line shape is consistent with that observed in a dye decolorizing peroxidase and attributed to a tryptophan radical.^18^

**Figure 2.**
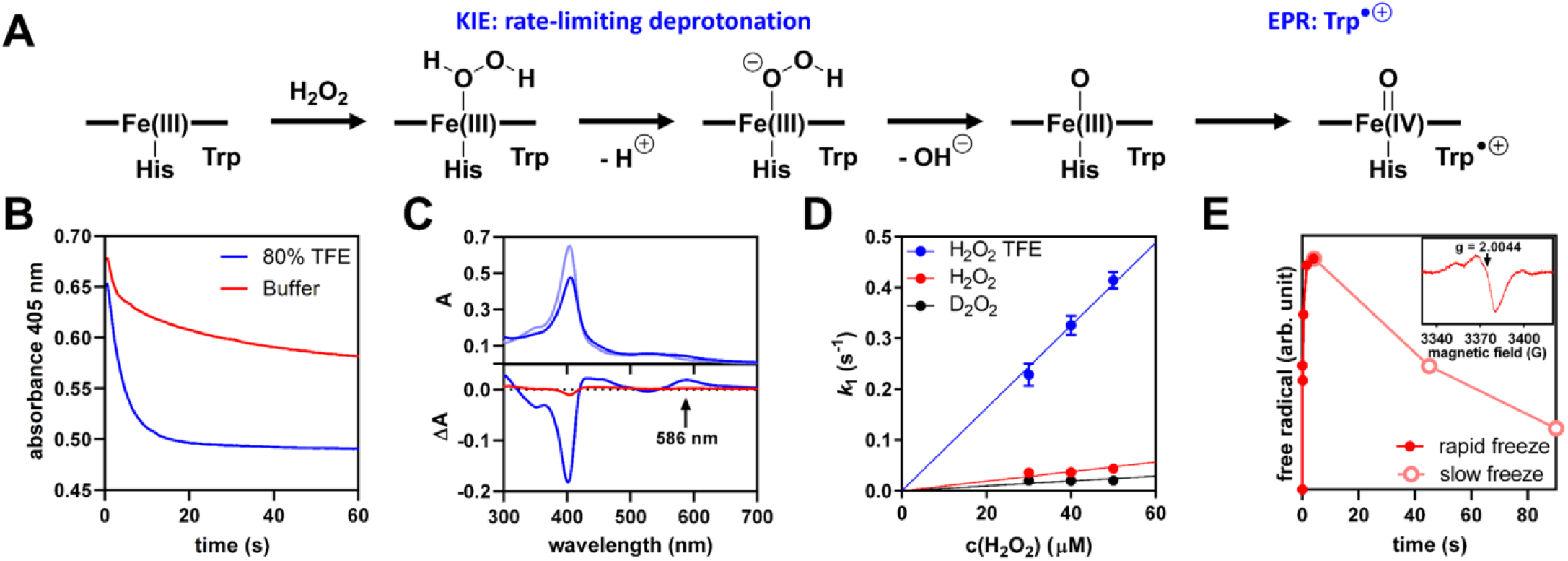
Accelerated formation and increased stability of Compound I in C45 in the presence of TFE. **A**, The proposed catalytic mechanism for formation of Compound I in C45. **B**, Reaction of 5 *μ*M C45 with 50 *μ*M H_2_O_2_ results in biphasic kinetics reflecting Compound I formation and heme degradation (buffer: blue, 80%TFE: red). TFE accelerates formation and slows down degradation. **C** Spectra in 80%TFE at 0.06 s and 60 s (top), and corresponding difference spectra in 80% TFE (blue) and buffer (red). Compound I has a Q band at 586 nm, that becomes most pronounced in 80% TFE at high temperatures (B and C recorded at 60 °C). **D** Rates of intermediate formation determined in 80%TFE (blue), buffer (red) and in 80% D_2_O with 5 *μ*M C45 at 5 °C. **E** Free radical concentration in the C45 samples frozen slowly or on a freeze-quench apparatus different time after H_2_O_2_ addition. In the *inset* the EPR spectrum of the Trp radical intermediate (also see Figure S2B). *Conditions, 67*.*5 μ*M C45, 13 *μ*M H_2_O_2_, 100 mM KCl, 20 mM CHES, pH 8.6, 25 °C.

The rate of Compound I formation (*k*_H2O2_) can be quantified from the observed rate in the peroxide-dependence (Figure 2D). Notably, running the reaction in 80% D_2_O decreased *k*_H2O2_ from 940 ± 30 M^−1^s^−1^ to 490 ± 20 M^−1^s^−1^ (Figure S2). The resulting kinetic isotope effects of 1.9 implies that the rate-limiting step in Compound I formation is the cleavage of a proton that can be exchanged with solvent, which probably represents the initial deprotonation of hydrogen peroxide at the distal heme face. These observations are in good agreement with the high *K*_M_ for H_2_O_2_ and may be related to the lack of distal histidine residues in C45, which are commonly employed in natural peroxidases to perform this deprotonation and to facilitate proton shuttling.

From Figure 2D, the addition of TFE gave rise to a notable increase in *k*_H2O2_, and a significant reduction in the rate of decay of Compound I in the absence of ABTS. TFE increases *k*_H2O2_ 9-fold compared to the reaction in buffer to 8130 ± 80 M^−1^s^−1^, mirroring the increased ABTS oxidation activity. Notably, addition of TFE also decreased heme degradation by up to 2-fold at 5 °C. The stabilization of the reactive Compound I intermediate allows C45 to maintain activity over a prolonged time, which directly relates to the increase in total turnover number observed for ABTS oxidation. Intriguingly, we observed the highest accumulation of Compound I at elevated temperatures (60 °C) in TFE. Under these conditions, a new Q band (586 nm) is clearly observable. Our kinetic data therefore indicate that the rate of Compound I formation, as well as its stability, is improved through conformational restriction of C45 by TFE. The unexpectedly high stability of Compound I at high temperatures may furthermore point to changes in temperature dependence arising from thermodynamic perturbation of the system.

### Structural and dynamical effects of TFE

TFE is known to stabilise α-helical folds by strengthening intramolecular hydrogen bonds and hydrophobic interactions,^19–20^ and appears to drive the adoption of a more active catalytic geometry in C45 (Figure 1). We therefore hypothesized that, in the absence of TFE induced conformational change, the rigidification of C45 might be the driver of changes in Compound I formation and stability that give rise to the enhanced *k*_cat_ values. We employed a range of spectroscopic methods to probe the putative restriction of conformational dynamics of C45 upon addition of TFE.

Far-UV CD spectra (Figure 3A) suggest an enhanced helical structure content in the presence of TFE, with a more negative ellipticity value at 222 nm (Θ_222_ nm). Thermal melts monitoring Θ_222_ nm in the absence of TFE (Figure 3B; red line) show a progressive increase in Θ_222_ nm, with an upward curvature that is indicative of relatively uncooperative unfolding of the helical structure. In the presence of TFE (Figure 3B; blue line) we only observe a linear increase in Θ_222_ nm with almost no curvature. That is, we do not observe the beginning of an unfolding transition as observed in the absence of TFE. These data track with our previous work in showing that the secondary structure of C45 is extremely thermally stable^7^ but that this stability is enhanced in the presence of TFE. Moreover, these data are consistent with our kinetic progress curves (Figure S1) in suggesting that TFE stabilises C45 with respect to denaturation.

**Figure 3.**
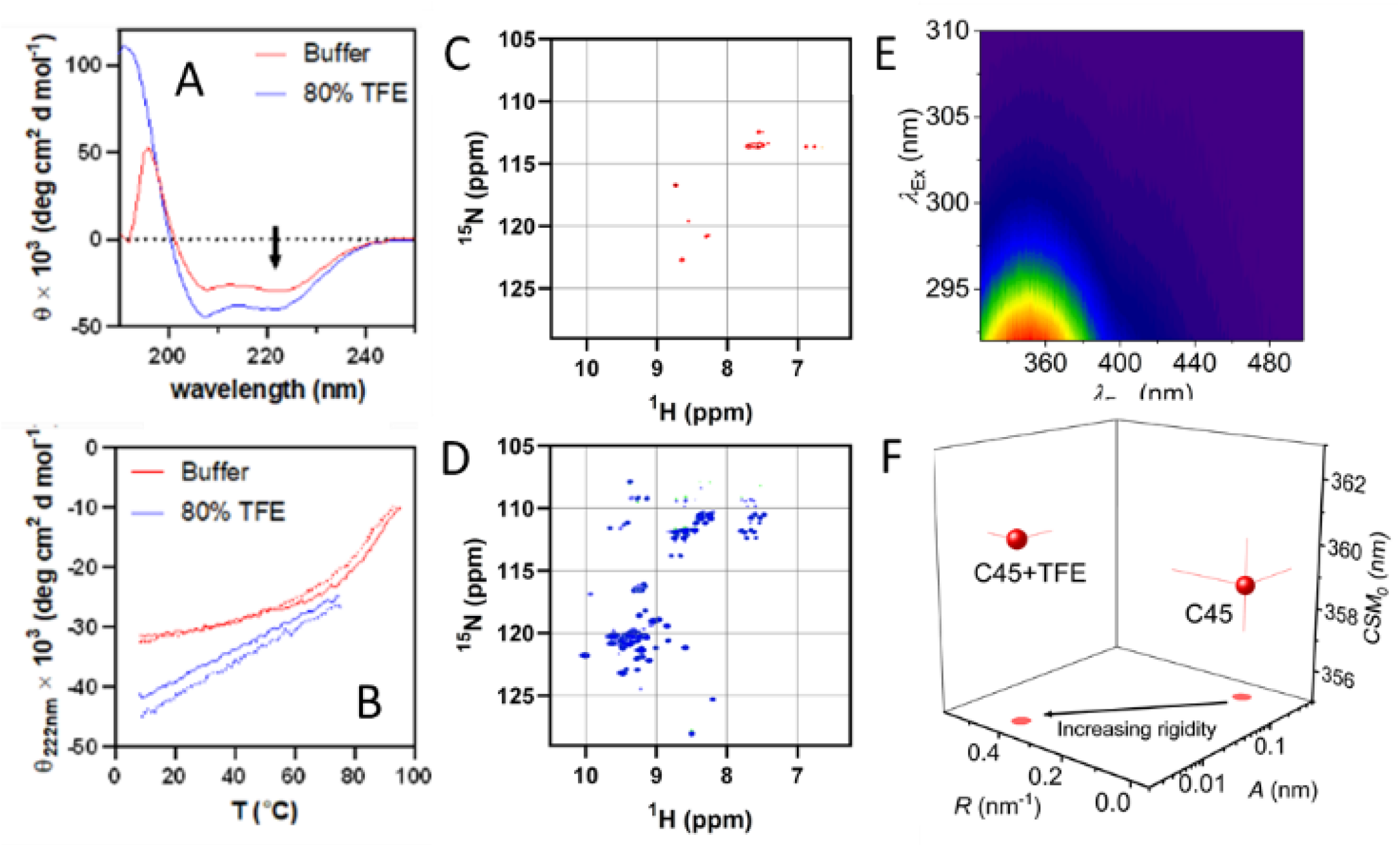
C45 rigidification in TFE. **A**, CD spectra and **B**, melting curves of C45 in buffer (red) and 80% TFE (blue). The dotted lines show the reversible refolding. *Conditions*, 5 – 10 *µ*M C45, 100 mM KCl, 20 mM CHES, pH 8.6, **C** and **D**, ^1^H -^15^N TROSY HSQC NMR spectra of C45 in buffer (panel C; red) and in 80% TFE (panel D; blue) *Conditions*, 250 *µ*M C45, 100 mM KCl, 20 mM CHES, pH 8.6, 10% D_2_O **E**, Example contour plot showing the change in structure of the emission spectra with increasing λ_Ex_. **F**, The QUBES data for C45 in buffer and in 50% TFE resulting from the fits shown in Figure S4. The solid black arrow indicates a decrease in the ratio *A*/*R*; interpreted as an increase in protein rigidity. *Conditions*, 4 *µ*M C45 in 50 mM HEPES buffer, pH 6.5, 15°C.

NMR spectroscopy allowed us to monitor the putative stabilizing effect of TFE at the residue level. The 2-dimensional ^1^H-^15^N TROSY-HSQC spectrum of C45 recorded in the absence of TFE (Figure 3C and S3) demonstrate relatively poor signal dispersion and resolution, suggesting a high degree of conformational flexibility, consistent with our previously reported C45 spectra and common to *de novo* designed helical bundles.^7,21–22^ In contrast, the addition of TFE to C45 results in a ^1^H-^15^N TROSY-HSQC spectrum (Figure 3D) with a notably increased number of sharper peaks. These data indicate that not only does C45 become more stable but is also less flexible/more compact. Rigidification slows down T2 relaxation, which increases intensity and sharpens peaks. The increased peak dispersion signals that C45 exists in a more defined conformational ensemble in TFE.

Far-UV CD cannot directly capture changes in protein flexibility and the protein concentrations used for NMR are high, compared to our kinetic studies, which potentially can affect molecular flexibility *via* macromolecular crowding.^23^ We therefore turned to our recent work using the red-edge excitation-shift (REES) phenomenon to further explore the change in C45 flexibility induced by TFE. Briefly, the REES phenomenon is observed as the inhomogeneous broadening of emission spectra with a decrease in excitation energy. This broadening reflects the presence of discrete solvent-solute interaction energies that are increasingly photoselected for by the change in excitation energy.^24^ We have demonstrated that this effect is capable of tracking changes in protein conformational sampling by monitoring intrinsic tryptophan emission, *via* a simple numerical model of the underlying REES effect.^25–27^ Indeed, we have demonstrated that this approach can identify differences in conformational sampling even between essentially identical crystal structures.^22^ We term our quantification and interpretation of the REES effect, QUBES (Quantitative Understanding of Bimolecular Edge Shift). We track changes in the broadening of Trp emission spectra as the change in the centre of spectral mass (CSM; See methods),

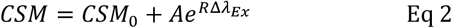

where the amplitude and curvature of the exponential is described by *A* and *R* values, respectively *CSM*_*0*_is the CSM value independent of *λ*_*Ex*_, the excitation wavelength andΔ*λ*_*Ex*_ is the change in excitation wavelength from 292 nm. *CSM*_0_ is then analogous to the use of Trp emission spectra to track changes in solvent exposure and infer tertiary structural changes i.e. as an increase in emission wavelength on unfolding (solvent exposure).^28^ That is, an increase in *CSM*_0_ would imply unfolding of the protein; increased solvent exposure of Trp. We have previously found that, for an invariant CSM_0_ value, an increased *A*/*R* value reflects a broader population of conformational states.^26–27^

Figure 3E shows the example raw REES data as the matrix of emission spectra *versus* changes in excitation energy. Plots of CSM *versus* excitation energy are shown in Figure S4, with the resulting parameters from fits to Eq 2 shown in Figure 3F. We note our tryptophan REES data are free from convolution with any iron-free heme as described in Methods and shown Figure S4. From Figure 3F, the *CSM*_0_ is the same within error and the *A*/*R* value in the presence of TFE is significantly smaller compared the absence of TFE. These data therefore suggest that the presence of TFE narrows the distribution of conformational states of C45. Potentially the narrower distribution of conformational states is the cause of the C45 rigidification in TFE. Moreover, our data do not suggest a measurable change in overall structure, since the extracted *CSM*_0_ values are essentially identical in the presence and absence of TFE (Figure 3F).

### Thermodynamic effects of altered C45 flexibility

To understand how the apparent protein rigidification affects the thermodynamic drivers of catalysis, we analysed the temperature-dependence of C45 steady-state turnover (Figure 4 and Table 1). From our previous studies, we do not expect strong binding of ABTS to C45.^7^ We have therefore conducted our steady-state kinetic analysis at the highest ABTS concentrations practically achievable. From Figure 4A and 4B, at all temperatures studied and both in the presence and absence of TFE, the steady-state kinetic data show a sigmoidal relationship, implying non-Michaelis-Menten type kinetics. These data contrast our work with C45 conducted at lower ABTS/H_2_O_2_ concentrations and or a different buffer system, suggesting the elevated substrate concentrations and or the change in buffer system have exposed a kinetic relationship that was not otherwise readily detectable. We note that in order to compare the kinetics of C45 to a natural enzyme (see below) we use a different buffer system with a much lower pH compared to previous reports ^7^ and so we do not expect the absolute magnitude of the extracted kinetics to be identical.

**Table 1.** Kinetic and thermodynamic parameters extracted from steady-state kinetics for both C45 and HRP.

|  | C45 | C45 + 50% TFE | HRP |
| --- | --- | --- | --- |
| $k_{\text{cat H}_2\text{O}_2}$ (S<sup>-1</sup>) | $13.5 \pm 2.4$ | $28.8 \pm 2.1$ | $638.0 \pm 45$ |
| $K_M \text{H}_2\text{O}_2$ (mM) | $1.6 \pm 0.40$ | $1.2 \pm 0.10$ | $0.05 \pm 0.01$ |
| $n$ | $1.6 \pm 0.2$ | $2.1 \pm 0.3$ | - |
| $T_{\text{opt}}$ (K) | - | 325.5 | 321.2 |
| $\Delta C_p^\ddagger$ (kJ mol<sup>-1</sup> K<sup>-1</sup>) | $0.04 \pm 0.19$ | $-0.28 \pm 0.19$ | $-1.65 \pm 0.75^b$ |
| $\Delta H_{T_0}^\ddagger$ (kJ mol<sup>-1</sup>)<sup>a</sup> | $5.4 \pm 0.95$ | $3.0 \pm 0.90$ | $24.1 \pm 4.70$ |
| $\Delta S_{T_0}^\ddagger$ (kJ mol<sup>-1</sup> K<sup>-1</sup>)<sup>a</sup> | $1.1 \pm 0.01$ | $1.1 \pm 0.01$ | $1.2 \pm 0.02$ |

a, $T_0 = 303$ K.

**Figure 4.**
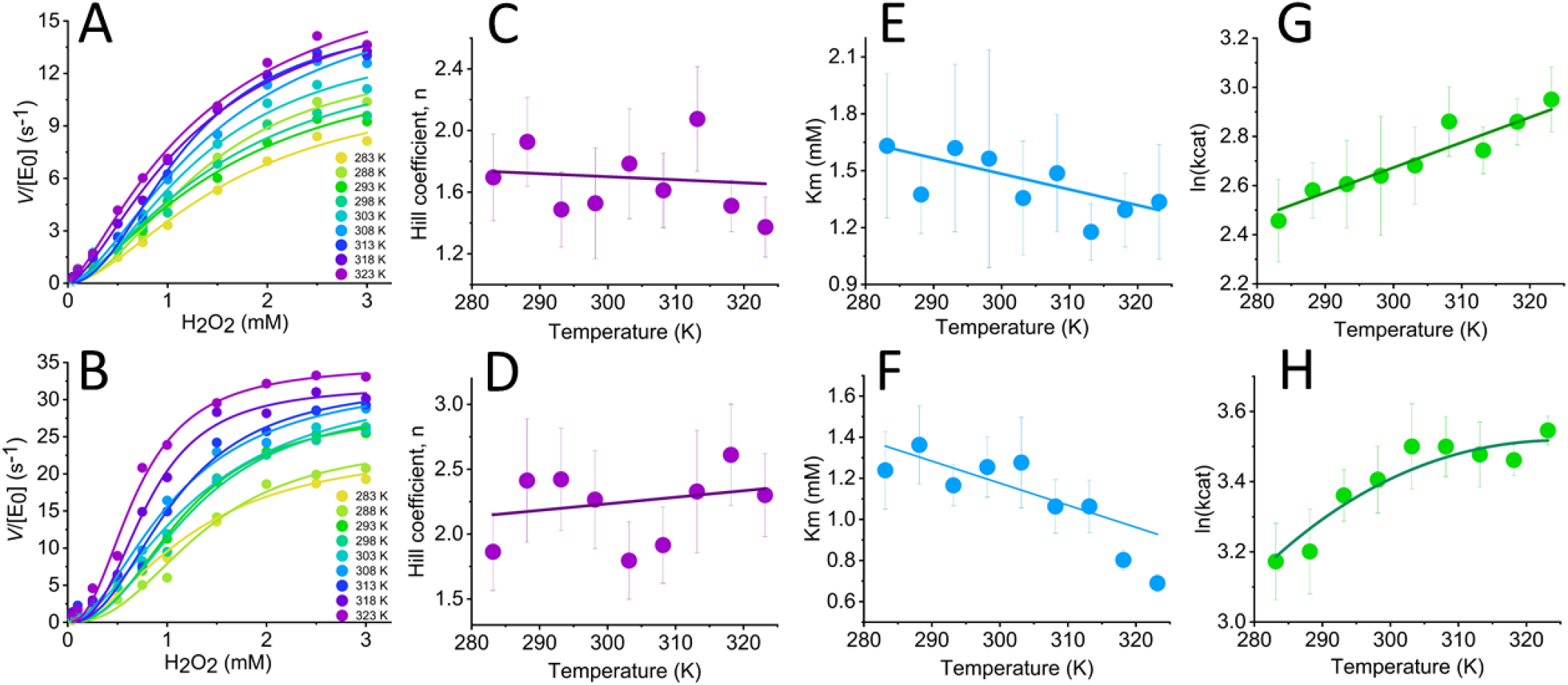
Temperature dependence of C45 turnover in the presence (at bottom) and absence (at top) of TFE. **A** and **B**, Concentration dependence of H_2_O_2_ versus rate of C45 turnover with increasing temperature (10-40 °C) in the absence (panel A) and presence of TFE (panel B). Solid lines are the fits of the data to Eq 3. **A**, in buffer and **B**, with 50% TFE (v/v). **B-H**, Temperature dependence of parameters extracted from steady-state data (panels A and B) for *n* (panels C and D), *K*_M_ (panels E and F) and *k*_cat_ (panels G and H). Solid lines in panels C-F are fits to a simple linear function and are to aid the eye only. Solid lines in panel G and H are the fit to Eq 1. Resulting parameters are given in Table 1. *Conditions*, 0.154 *µ*M C45 and 73 mM ABTS in 50 mM HEPES buffer, pH 6.5.

It is common^29^ to fit such apparently sigmoidal steady-state data to the Hill equation:

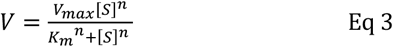

where the Hill coefficient, *n*, captures the deviation from hyperbolic Michaelis-Menten curves. The Hill coefficient is often attributed to allosteric binding effects.^30–31^ The magnitude of *n*, both in the absence and presence of TFE, signals positive cooperativity; *n* = 1.6 ± 0.2 and 2.1 ± 0.3 at 293 K, respectively, noting that these values are within error of each other. From Figure 4C and 4D, we find the magnitude of *n* is essentially invariant with temperature within the error of the measurement, in both the presence and absence of TFE. The extracted *K*_M_ values are again similar in the presence and absence of TFE (Figures 4E and 4F; Table 1); *K*_M_ = 1.6 ± 0.4 and 1.2 ± 0.1 mM, respectively. Potentially the *K*_M_ shows some decrease with increasing temperature, particularly in the presence of TFE, though we are reluctant to compare these putative trends given the magnitude of the calculated error. Finally, as with our findings above, the extracted *k*_cat_ is larger in the presence of TFE (Table 1).

We acknowledge that diagnosing true allosteric cooperativity is challenging, not least for a small artificial enzyme system where one of the substrates does not have an obvious formal binding site as we describe above. Without a structural underpinning for the effect we cannot consider the detailed putative allosteric mechanism. Instead, accurately tracking the steady-state kinetics of C45 allows us to extract robust kinetic data, from which we are able to determine key thermodynamic parameters from temperature dependence studies.

The temperature dependence of the extracted *k*_cat_ values are shown in Figure 4G and 4H in the presence and absence of TFE (50% v/v), respectively. The extracted parameters resulting from fits to Eq 1 are given in Table 1. From Figure 4G, 4H and Table 1, we find there is evident curvature in the temperature dependence plots in the presence of TFE only, manifesting as a shift from a ∼zero 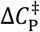 (0.04 ± 0.19 kJ mol^−1^ K-1) to a measurably negative 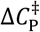 in the presence of TFE (−0.28 ± 0.19 kJ mol-1 K-1).

Given we have demonstrated that C45 is more stable with TFE present (above), the apparent negative 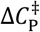 value in the presence of TFE would not appear to be due to local unfolding. Moreover, our stopped-flow and absorption data studies (discussed above) indicate that the chemical mechanism is essentially invariant. We acknowledge that we cannot rule out a change in rate limiting step. In the absence of such confounding factors, a negative 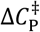 indicates a difference in the distribution of vibrational modes between the ground and transition state: In a simple sense, a difference in the selective rigidification between the ground and transition state ensemble. Based on previous work we anticipate this could involve larger scale motions throughout the protein as illustrated previously in other systems,15,32 though we do not rule out more localised effects.

The presence of TFE induces a negative 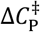, which manifests as curvature in temperature dependence of *k*cat. A corollary of this is the presence of a temperature optimum of the reaction (*T*opt) in the presence of TFE (*T*opt= 325.5 K). Given there is no measurable curvature in the temperature dependence data in the absence of TFE, we cannot assign a *T*opt. Our data therefore indicate that whilst TFE increases the thermal stability of C45, it also induces a *T*opt at a temperature much lower than unfolding (by comparisons to our CD data; Figure 3). Rigidifying the enzyme, induced by TFE, appears to decoupl*e* the stability of C45 and the temperature optimum of catalysis.

### Comparison to the kinetics and thermodynamics of a natural peroxidase

We wish to explore the effect of a globally different relationship of protein conformational dynamics to enzyme turnover, beyond what is accessible with our TFE studies. To that end we compare C45 to a natural enzyme that catalyses a similar chemical reaction, albeit with a more typical (larger) protein structure. Horseradish peroxidase (HRP) is an excellent model system for this purpose. HRP is ∼44 kDa with a largely α-helical structure, with peroxidase activity mediated by a b-type heme cofactor.33

First we assessed the role, if any, of protein conformational dynamics on HRP turnover. Whilst HRP can potentially be stable to TFE,^34^ we find that in our buffer system this is not the case and the extracted rate constant decreases with increasing TFE (Figure S5), presumably due to HRP unfolding. Instead of using TFE as with C45, we were able to perform combined temperature and pressure studies that provide more detailed insights into the conformational state equilibria affecting catalysis.

Non-denaturing hydrostatic pressure is an established method for demonstrating the sensitivity or not of enzyme turnover to changes in the equilibrium of protein conformational states.^35–38^ As a broad framework, one expects a significant pressure dependence on enzyme activity, in cases where the turnover is affected by the proteins conformational dynamics.^39–41^ We have previously demonstrated a correlation of a pressure-dependent 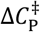 with differences in global protein flexibility.^26^ Figure 5A and 5B show the combined pressure-temperature dependence of *k*_cat_ for HRP and the resulting pressure dependence of the values extracted by fitting to Eq 1 and given in Table 1. Example steady-state Michaelis-Menten data are shown in Figure S6. We note that our progress curves are linear at elevated pressures (Figure S7), and our data are fully reversible with pressure, showing that our data are not convolved with unfolding at the pressures we use.

**Figure 5.**
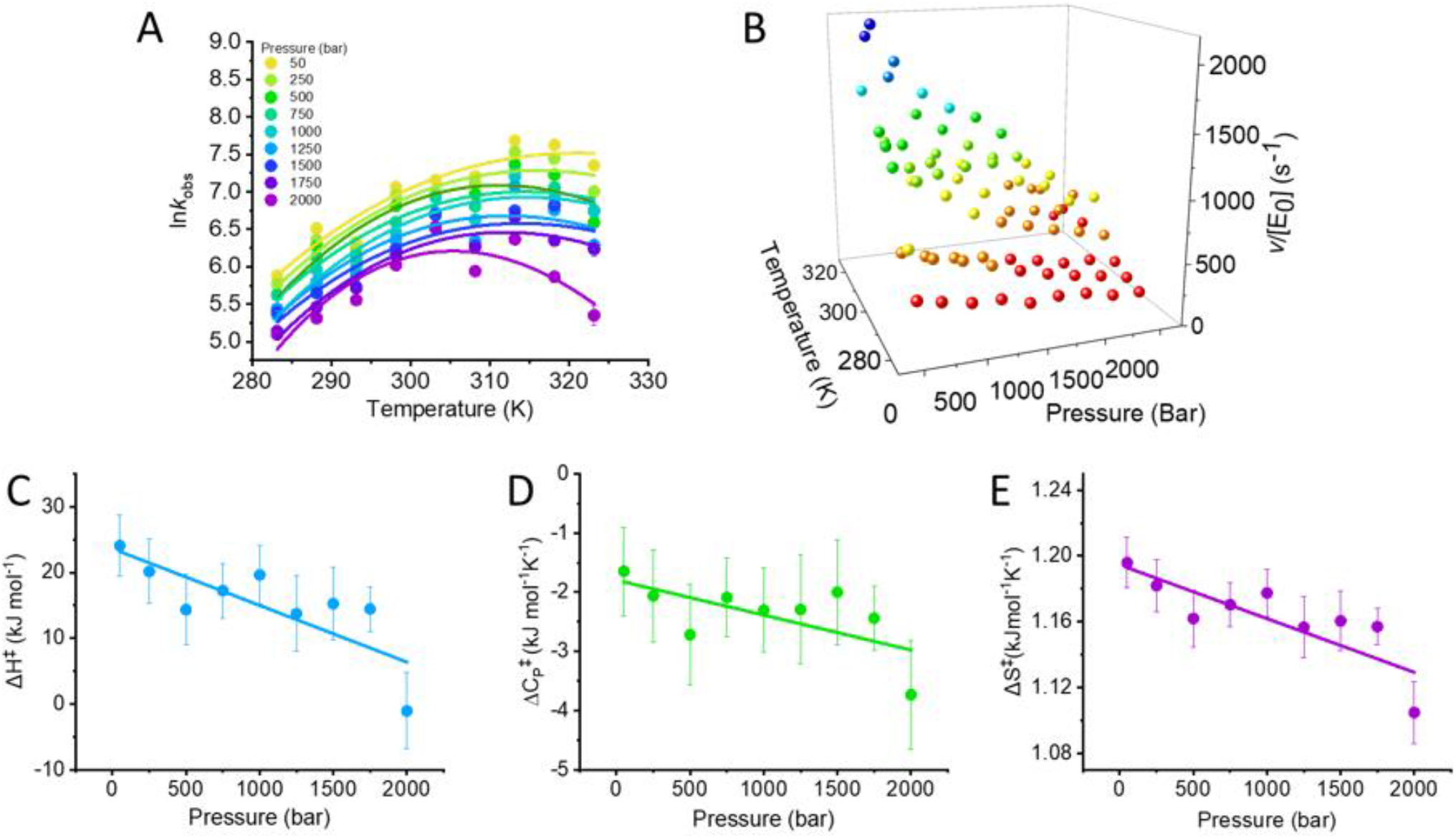
Combined pressure and temperature dependence of HRP turnover (panels **A** and **B**). The solid lines in panel **A** are the fits to MMRT Eq 1. **C-E**, Pressure dependence of resulting parameters from fits in panel **A**. The solid lines are to aid the eye only and illustrate the prevailing trend in the data with the resulting parameters plotted in panels **C**-**E**. Resulting parameters are given in Table 1. *Conditions*, 0.2 nM HRP, 1.8 mM H_2_O_2_ and 73 mM ABTS in 50 mM HEPES buffer, pH 6.5.

From Figure 5A and 5B, the *k*_cat_ of HRP varies significantly with both pressure and temperature. That is, we observe a pressure-dependent change in *k*_cat_ at all temperatures studied, suggesting that altering the distribution of conformational states, impacts on HRP turnover, and in turn implying a role for HRPs conformational dynamics in enzyme turnover. We note that the substrates in the absence of enzyme, show essentially no chemical turnover on the timescales of our assays, and so the pressure dependence we observe is due to the enzyme.

Our pressure/temperature matrix (Figure 5A and 5B) allows us to further explore whether the thermodynamics associated with enzyme turnover are pressure dependent, and to more specifically infer the role of protein conformational dynamics. Figures 5C-D show the pressure dependence of the extract values from Eq 1 at each pressure studied. From these data it is clear all thermodynamic parameters are pressure dependent, showing an approximately linear decrease with respect to pressure. It is interesting to note that the decrease in 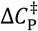 is also correlated with a general decrease in *T*_opt_ (Figure 5A), consistent with the predictions from MMRT. These data therefore suggest that altering the equilibrium of protein conformational states in HRP is sufficient to alter the thermodynamics of enzyme turnover, analogous to the effect of TFE on C45.

From Table 1, the kinetics and thermodynamics of C45 turn-over in the presence of TFE approach those of HRP; with an elevated *k*_cat_, a measurably negative 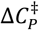 and a similar *T*_opt_. It is therefore tempting to speculate that the rigidification of C45 has given rise to a more ‘natural’-like enzyme at least in terms of the thermodynamics of enzyme turnover. This suggestion is satisfying because one anticipates enzyme active sites are significantly organised by the bulk protein.

## Conclusions

*De novo* enzymes have enormous potential to be platform biocatalysts, with significant scope for engineering and tuning activity. It is common for enzyme engineering efforts with ‘natural’ enzymes to attempt to rigidify the overall structure/active site, primarily for enhanced thermal stability, but also to precisely engineer specific active site geometries. Here, we use a model peroxidase with excellent catalytic activity to explore the effect of altering the rigidity of an artificial enzyme on protein dynamics, stability and catalytic activity. By using a simple shift in solvent system we are able to tune the rigidity of C45, increasing thermal stability but also enhancing the rate of turnover. The combined effect of increased stability and activity leads to an enzyme with a significantly increased total turnover number compared to the parent system. Potentially, rigidification of C45 drives the adoption of a more active state, *via* e.g. stabilisation of Compound I as we discuss above. That a simple solvent change is able to increase the TTN of the *de novo* enzyme by over 30 fold points to a potentially simple inexpensive route to tune such enzymes. We note that the link between protein flexibility/rigidity and activity will be system dependent, but smaller, less complex systems like C45 provide a window into this relationship.

Our thermodynamic studies point to a most intriguing finding, namely that increasing the rigidity of C45 decouples thermal stability from the temperature optimum of reaction. That is, for C45 we observe global rigidification of the protein at large, but simultaneously a difference in the distribution of vibrational modes between the ground and transition state, giving rise to a negative 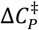 and a measurable *T*_opt_, well below a measurable unfolding transition. C45 is therefore a key case study in illustrating that the anticipated relationship between stability, activity and response to temperature does not always track in the expected ‘text-book’ way. Moreover, it is notable that these findings are with a small, *de novo* enzyme *versus* a large natural enzyme. This study adds to the growing evidence that a negative activation heat capacity is indicative of ‘enzyme like’ behaviour, and can correlate with catalytic activity.^42^

Comparison of the thermodynamics of reaction between C45 and a natural enzyme (HRP) show that for both proteins, perturbing the equilibrium of conformational states can alter the thermodynamics of catalysis and particularly the magnitude of 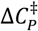. This finding is all the more interesting since the rigidification appears to tune the thermodynamics of C45 to be more similar to a larger ‘natural’ enzyme, which catalyses a similar chemical reaction. Our data then provoke important questions around the ‘optimal’ flexibility of an enzyme and the role of the protein scaffold at sites distal to the immediate active site volume.^13^

## Materials and Methods

### Protein production and purification

C45 was produced and purified as previously described. Briefly, C45 was expressed in T7 express *E. coli* BL21 (DE3) co-transformed with vector pEC86 encoding for the c-type cytochrome maturation system Ccm. Protein expression was induced with the addition of isopropyl β-D-1-thiogalactopyranoside (IPTG). A cell pellet was collected and lysed, then purified using nickel affinity column chromatography. The 6-histidine tag was cleaved using tobacco etch virus protease (100 *µ*g/L expression culture) under anaerobic conditions in the presence of a reducing agent (1 mM Tris(2-carboxyethyl)phosphine). The cleaved protein was further purified using nickel affinity column chromatography and size exclusion chromatography, flash frozen using liquid nitrogen and stored at −70 °C. HRP was purchased from Sigma-Aldrich.

### CD spectroscopy

CD experiments were carried out on a Jasco J-810 spectrophotometer under nitrogen atmosphere, using a sealed quartz cuvette. A baseline of the buffer solution including cosolvent was obtained and subtracted from protein measurements. Scans were taken at a rate of 100 nm per minute and a temperature of 25 °C. Thermal denaturation studies were conducted with a ramp of 1 °C min^−1^, starting at 8 °C.

### REES spectroscopy

All fluorescence measurements were performed using a Perkin Elmer LS50B Luminescence Spectrometer (Perkin Elmer, Waltham, MA, USA) connected to a circulating water bath for temperature regulation (1 °C). Samples were thermally equilibrated by incubation for 5 minutes at the given conditions prior to recording measurements. For all samples, the corresponding buffer control was subtracted from the spectra for each experimental condition and this also removed the Raman water peak.

The fluorescence emission spectra are typical of Trp fluorescence data, with the exception that an additional minor spectral feature at ∼440 nm that can be attributed to free heme. The quantification of the REES data relies on accurate extraction changes to the emission spectra and so the additional band would convolve the measurement. We have therefore numerically modelled each of the spectra using a sum of two skewed Gaussians;

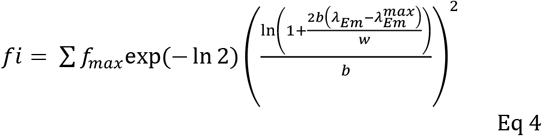

where *f*_i_ is the measured fluorescence intensity, *f*_max_ is the maximum emission intensity at wavelength, with a full width at half maximal of *w* and the ‘skewness’ is controlled by *b*. Fluorescence spectra can be accurately modelled and deconvolved by fitting to such functions. By fitting to a sum of two skewed Gaussians we are able to accurately model the spectral component attributable to Trp emission alone (see Figure S4). We use the resulting model attributable to Trp emission, deconvolved from any contribution from iron-free heme to extract the value of CSM,

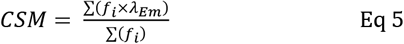

The CSM can then be calculated more accurately. Data collected in a thermostated cell holder (15 °C). Excitation and emission slit widths were 4 nm and C45 tryptophan emission was monitored from 325 to 550 nm. The excitation wavelength was subsequently increased in 1 nm steps for a total of 19 scans. C45 was present at a concentration of 4 *µ*M, with some conditions containing 50% TFE (v/v). Data fitting and plotting was performed using ORIGINPRO 2019, 9.6.0.172 (MicroCal, Malvern, UK).

### NMR spectroscopy

Experiments were performed on a Bruker Avance III HD 700 MHz instrument equipped with a 1.7mm TXI Z-gradient probe at 298 K. NMR samples contained 250 *μ*M ^15^ N-labeled C45 in buffer (20 mM CHES, 100 mM KCl, pH 8.6) containing 10 % D_2_ O. A second sample was prepared with 80 % deuterated TFE, 10% buffer and 10% D_2_O.

### EPR spectroscopy

Low temperature EPR spectra were recorded on a Bruker EMX (X-band) EPR spectrometer with the use of an Oxford Instruments liquid-helium system and a spherical high-quality ER 4122 (SP 9703) Bruker resonator. The instrumental conditions used to record the EPR spectra were as follows: microwave frequency ν_MW_ = 9.4677 GHz, microwave power P_MW_ = 0.79 mW, modulation frequency ν_m_ = 100 kHz, modulation amplitude A_m_ = 3 G, time constant τ= 82 ms, scan rate V = 0.60 G/s, number of scans per spectrum NS = 4. The slow freeze samples were prepared by immerging an EPR tube with reacting mixture to methanol kept on dry ice. Rapid Freeze-Quenched (RFQ) EPR samples were prepared on an isopentane-free apparatus as described previously.^43^

### Steady-state kinetics

All reactants were pre-incubated in the experimental concentration of cosolvent and mixed immediately prior to measurement. Baseline measurements were subtracted from results. Kinetics data were either collected by plate reader (Synergy neo2 multi-mode reader by BioTek Instruments using StarLab 96-well microplates with flat-bottomed, round wells), or using a UV/Vis spectrophotometer (Agilent Cary 60 UV-Vis spectrometer) fitted with temperature regulation, in either 1 cm/3 mm/1 mm cuvettes, or mounted in a high-pressure cell (see below). All experiments for both enzymes were performed in 50mM HEPES, pH 6.5 unless stated otherwise.

In all cases, reactions were initiated by the addition of either C45 or HRP and the formation of the ABTS radical cation was monitored over a range of wavelengths from 414 nm -465 nm for which the molar extinction coefficients were calculated. The C45 temperature dependence kinetics was measured over a temperature range of 283.15 K – 323.15 K in 5 K increments.

### Pressure dependence studies

An ISS high-pressure cell (ISS, Champaign, UL, USA), fitted with a custom mounting to an absorbance spectrometer connected to a circulating water bath for temperature regulation, was used to record all pressure measurements. In all cases, reactions were initiated by the addition of HRP (0.02 nm) and the formation of the ABTS radical cation was monitored using wavelengths between 414 nm -465 nm for which the molar coefficients were calculated. The experiment was carried out over a temperature range of 283.15 K – 323.15 K in 5 K increments and a pressure range of 50 Bar – 2000 bar over 9 increments.

### Pre-steady-state kinetics

Formation of Compound I was measured by mixing C45 with H^2^O_2_ in an Applied Photophysics SX20 Stopped-Flow spectrometer connected to a photodiode array detector. and was recorded on a log-time-scale. Data between 380 nm and 700 nm were fitted to a single exponential accounting for Compound I formation plus a linear function reflecting the decay. Data were fitted from 1 s to 20 s for 80% TFE, and 1 s to 120 s for the data without TFE. Data were fitted globally with the rate of Compound I formation (*k*_1_) shared between all wavelengths. For TFE, a substantial background signal associated to scattering of micelle rapture and formation during mixing was observed. The background signal was obtained by mixing C45 in 80% TFE without peroxide and subtracted from the reaction kinetics before fitting. Solvent isotope effects were determined by running the reaction in 80% D_2_O.

## ASSOCIATED CONTENT

Supporting information including kinetic progress curves, EPR and NMR data, raw REES data and steady state enzyme kinetic data. This material is available free of charge via the Internet at http://pubs.acs.org.

## AUTHOR INFORMATION

### Funding Sources

We acknowledge BBSRC funding for SH’s studentship as part of the SWBio doctoral training partnership. HAB thanks the Swiss National Science Foundation for a Postdoctoral Mobility fellowship. BF thanks the EPSRC-funded Bristol Centre for Functional Nanomaterials for her studentship (EP/G036780/1). AJM thanks EPSRC for funding (grant number EP/R027129/1). JLRA and AJM thank the BBSRC for funding (BB/R016445/1), and JLRA, CM and AJM thank BrisSynBio, a BBSRC/EPSRC Synthetic Biology Research Centre (Grant Number: BB/L01386X/1).

#### ABBREVIATIONS

REES: red edge excitation shift
MMRT: Macromolecular rate theory
TFE: 2,2,2-Trifluoroethanol, TFE
CSM: centre of spectral mass
QUBES: quantitative understanding of biomolecular edge shift

## Table of Contents

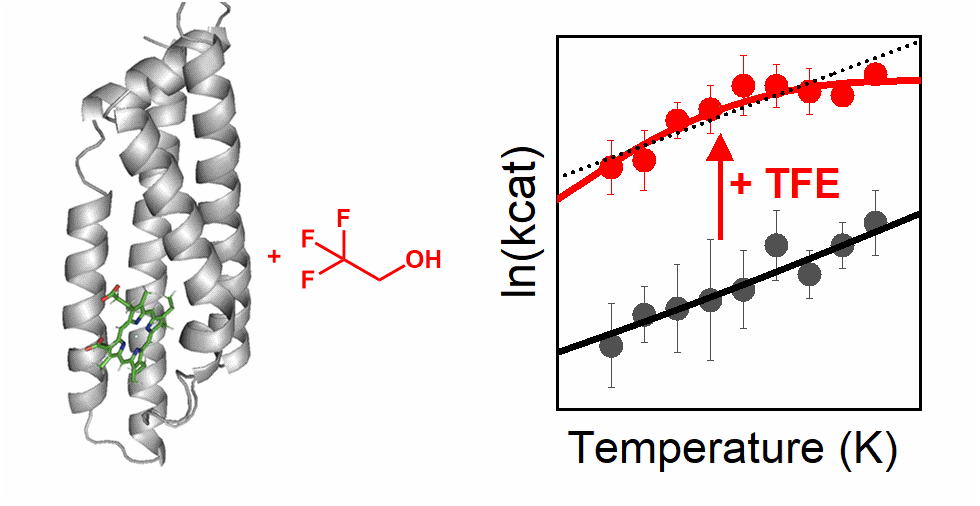

## Supporting Information

**Figure S1.**
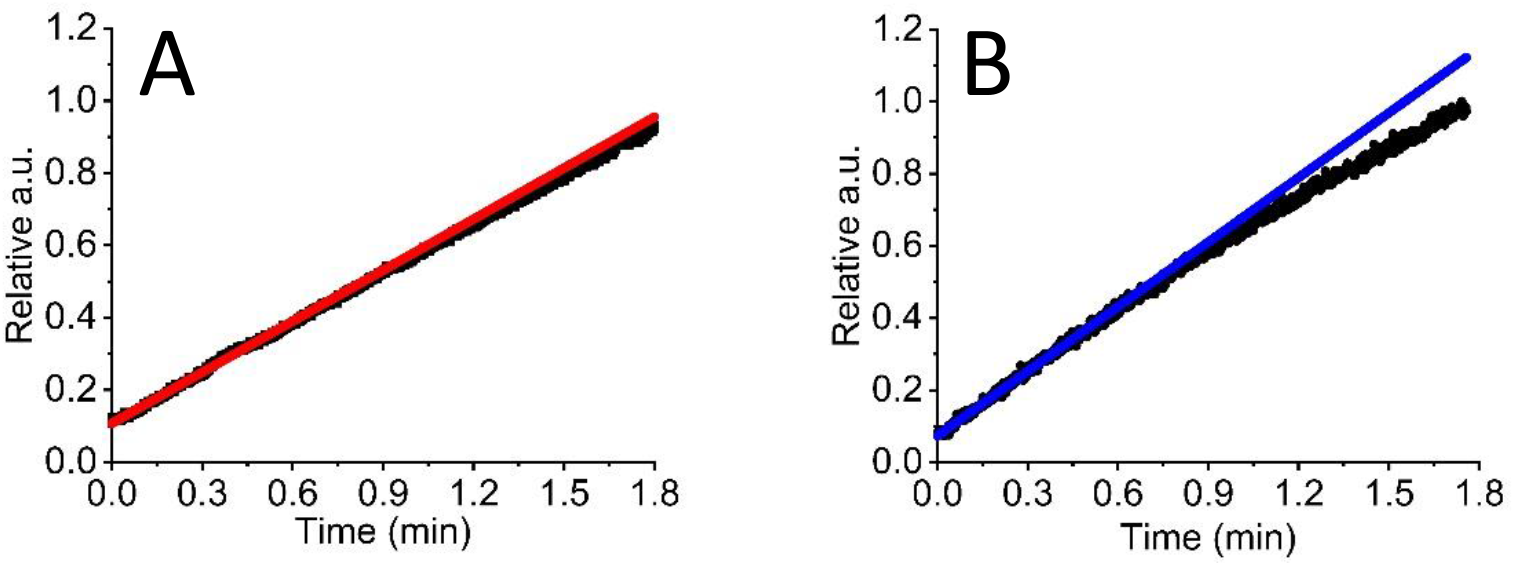
Kinetic progress curves showing the stabilising effect of TFE, H_2_O_2_ versus rate of C45 turnover in the presence (panel **A**) and absence of TFE (panel **B**). Solid lines are the fit of a simple linear function to the firs 0.5 min of data, showing the deviation from linearity (initial rate) with respect to time. *Conditions*, 0.15 *µ*M C45 and 73 mM ABTS in 50 mM HEPES buffer, pH 6.5 at 30 °C.

**Figure S2.**
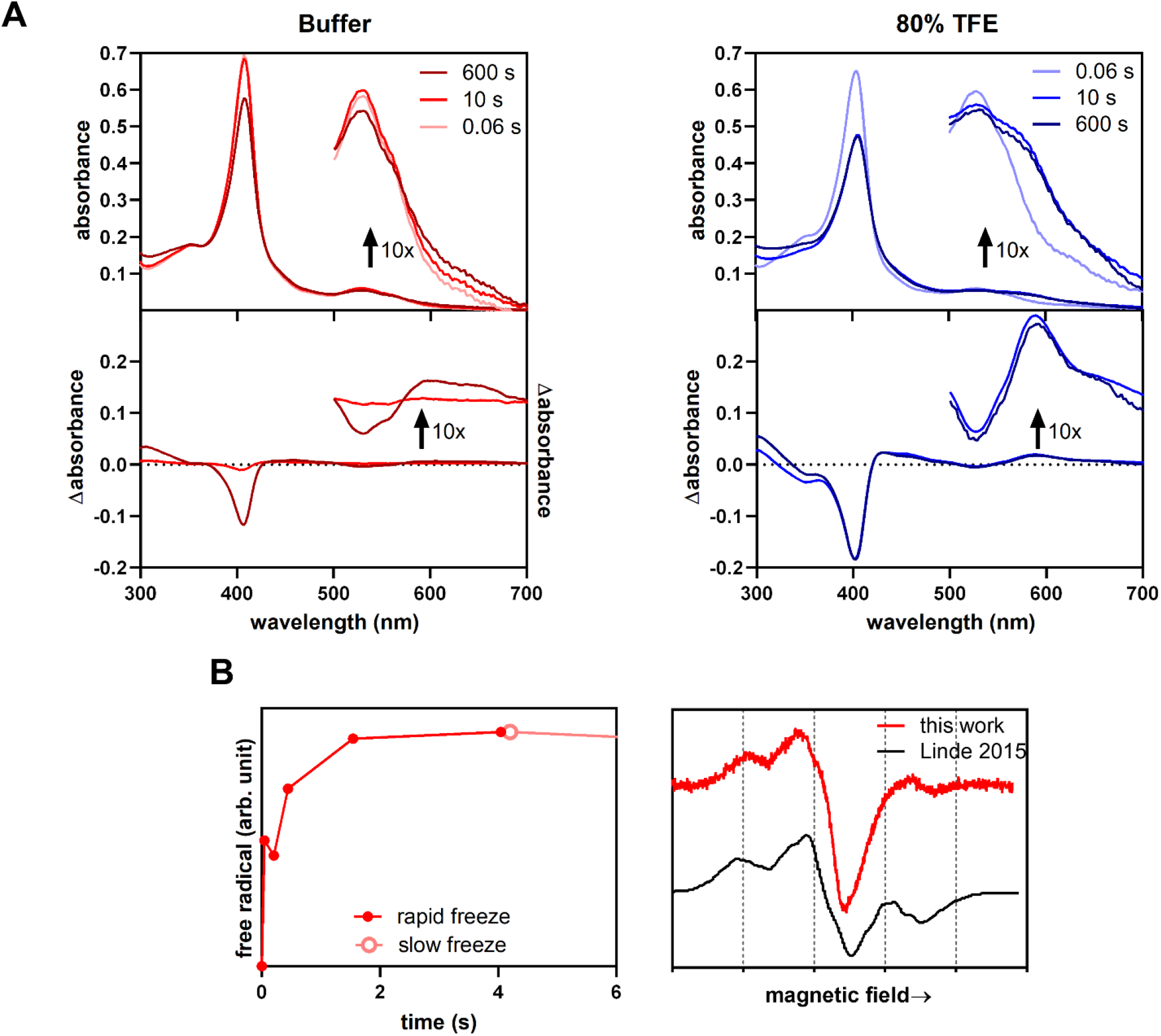
Monitoring Compound I formation. **(A)** Rapid mixing of peroxide and C45 in buffer (left, red) and 80% TFE(right, blue) leads to formation of Compound I, as indicated by the UV/Vis spectra (top). To illustrate spectral changes associated with the reaction, difference spectra compared to the first time point at 0.06 s are shown (bottom). More of the intermediate-associated Q-band at 586 nm is present after 10 s in TFE compared to buffer, signalling rapid formation of Compound I in the presence of the cosolvent. The Q-band and Soret peak degrade much slower in TFE compared to buffer, indicating that Compound I is more stable in the fluorinated solvent. **(B)** Detailed view of the initial part of the kinetic curve of the radicals presented in Figure 2E. The slow and rapidly freeze-quenched samples have different densities (and therefore different EPR signals intensities); the two sets of data have been brought to a common value at the common time point – 4 s. The unusual, three component EPR spectrum of C45 (red) is plotted on a common magnetic field scale with the published spectrum of a dye decolorizing peroxidase (DyP) assigned to a Trp radical.^1^ The spectrum of DyP from 21 (black) has been digitized using UN-SCAN-IT 6.0 (Silk Scientific). As the microwave frequencies used in ref 21 and in this study are slightly different, the EPR signals appear at slightly different values of the magnetic field (horizontal axis). Because of that, the values of the field, increasing from left to right, are not shown in the figure, but rather the vertical gridlines at a 20 Gauss intervals indicate the overall scale of the scan. The two spectra have been aligned by a common value of the g-factor.

**Figure S3.**
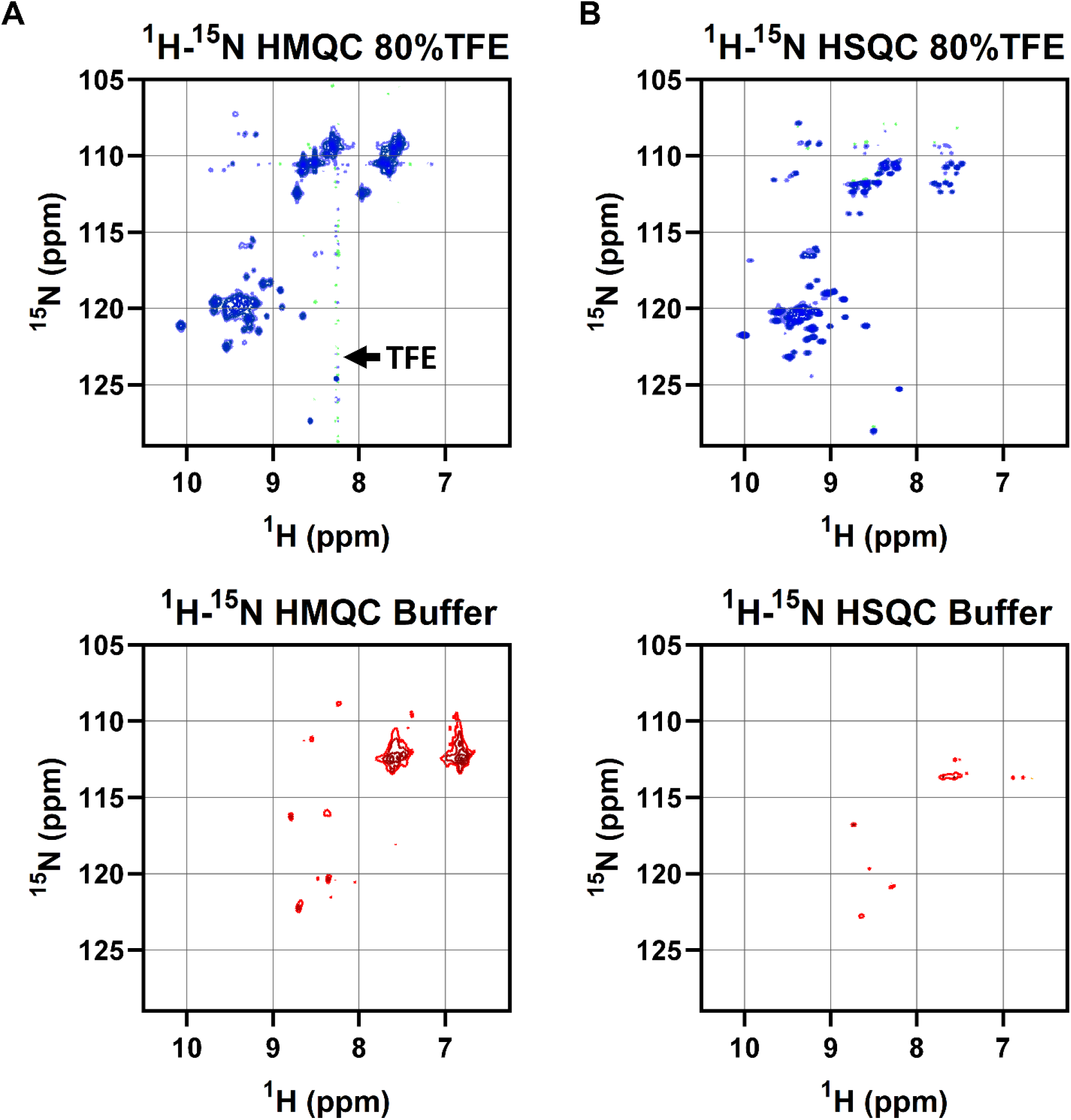
NMR spectra of C45 in 80% TFE (top, blue) and buffer (bottom, red) **A)** ^1^H-^15^N HMQC (left) and **C)** HSQC (right) spectra become more disperse and more peaks are apparent after addition of 80% TFE.

**Figure S4.**
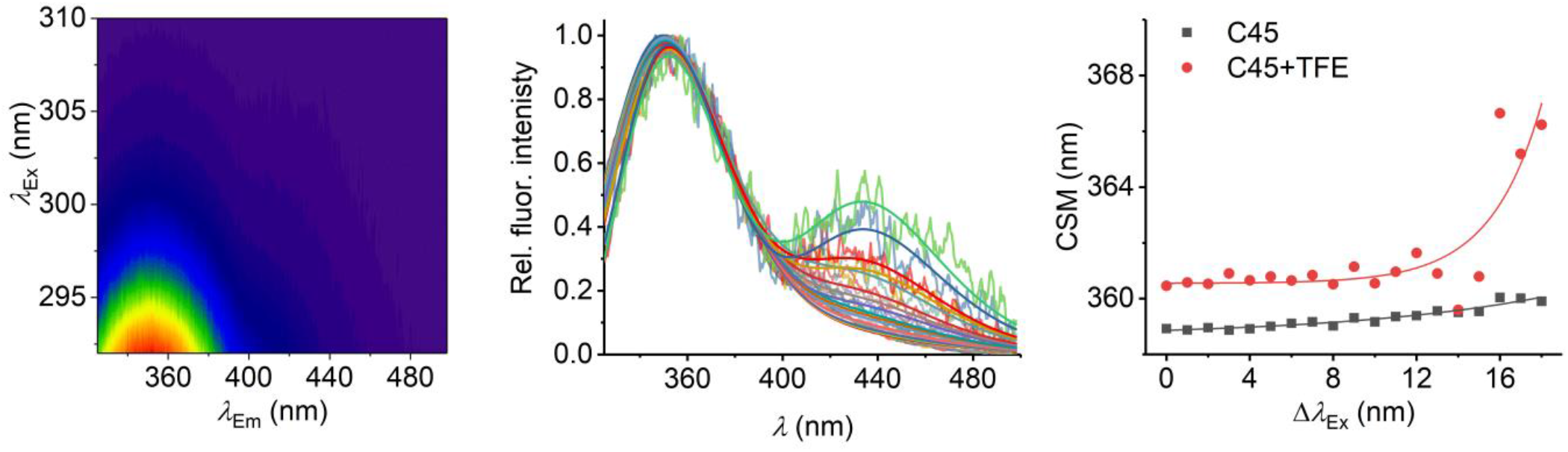
REES spectroscopy and extracting QUBES data. **A**, Raw REES data. **B**, Fitting of raw REES data in panel A using Eq4 as described in *Methods*. **C**, Plot of CSM *versus* excitation energy for C45 in the presence and absence of 50% TFE. Solid lines show fits to Eq 2. *Conditions*, 4 *µ*M C45 in 50 mM HEPES buffer, pH 6.5, 15°C.

**Figure S5.**
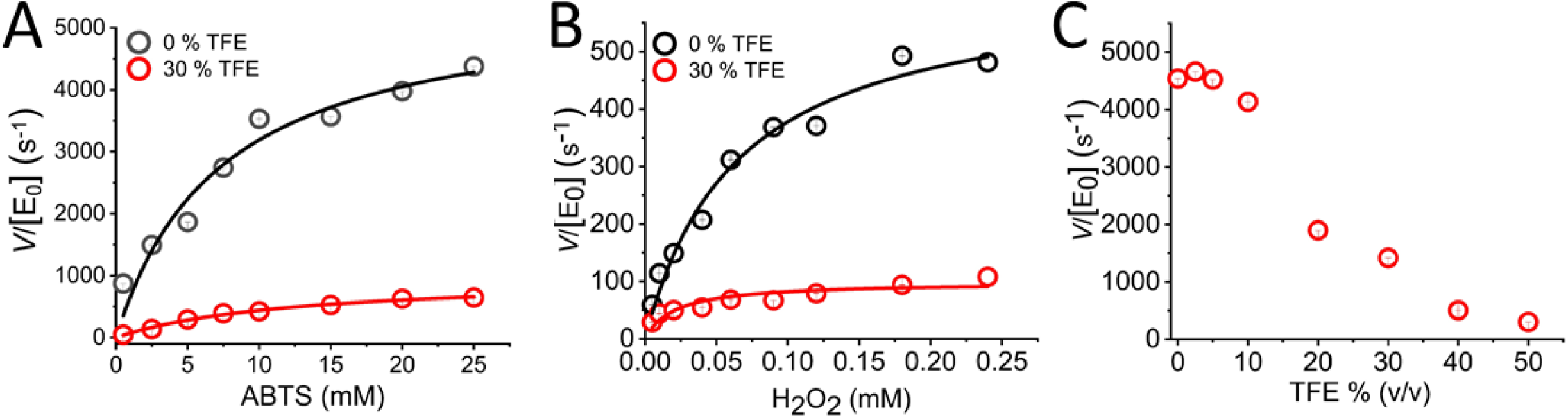
The steady state kinetics of HRP turnover at 0% and 30% TFE recorded over a range of ABTS and H_2_O_2_ concentrations (panel **A** and **B** respectively). **C**, Effect of increasing TFE % (v/v) on maximal rate of HRP, all data was recorded in triplicate and SE is shown. *Conditions*, Panel A 0.5 nM HRP, 0.5 mM H_2_O_2_, Panel B 0.2 nM HRP, 30 mM ABTS and panel C 0.2 nM HRP, 0.5 mM H_2_O_2_ and 30 mM ABTS, all in 50 mM HEPES buffer, pH 6.5.

**Figure S6.**
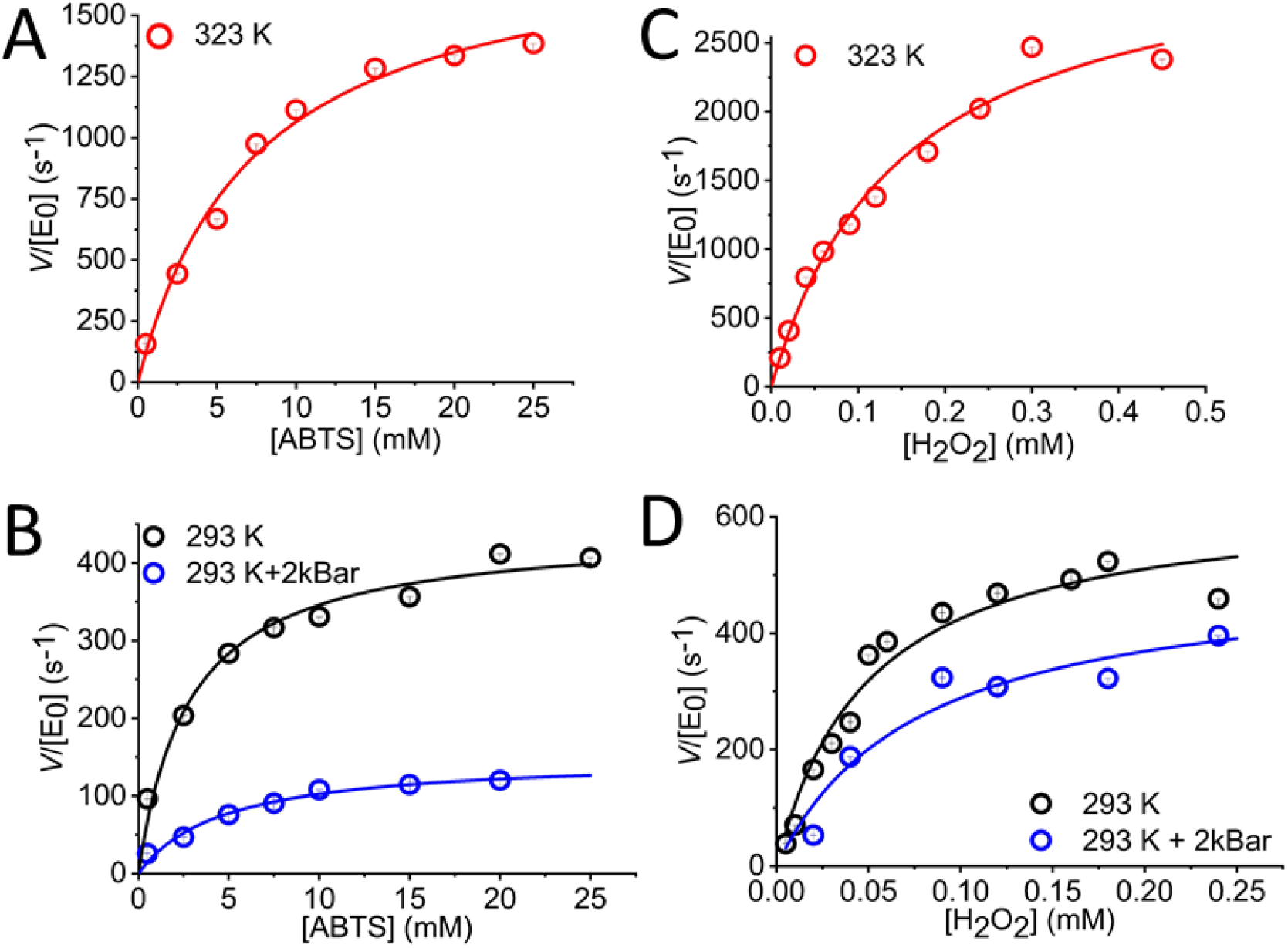
The steady state kinetics of HRP turnover varying [ABTS] (panel **A and B**) and [H_2_O_2_] (panel **C and D**), All data were recorded in triplicate and error bars represent the standard error. *Conditions*, 0.5 nM HRP, 0.13 mM H_2_O_2_ (panel A), 30 mM ABTS (panel B) in 50 mM HEPES buffer, pH 6.5.

**Figure S7.**
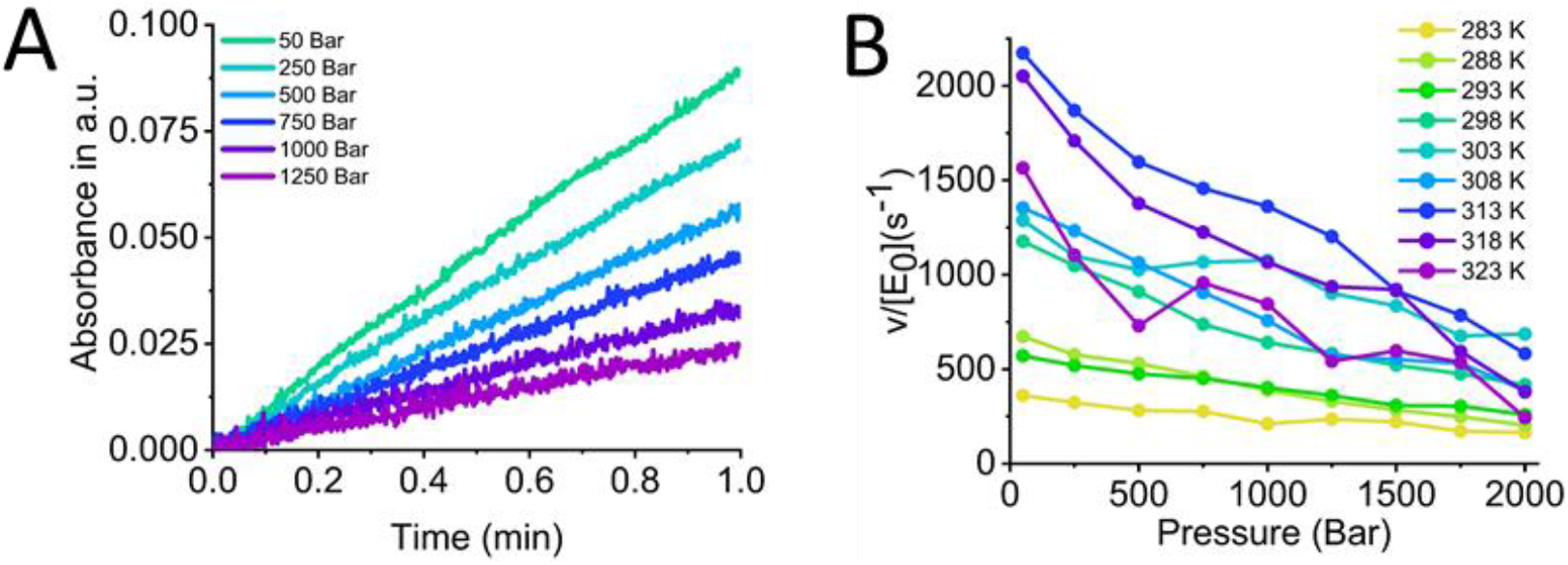
Combined pressure and temperature dependence of HRP turnover. **A**, Raw data shows HRP progress curves are linear at elevated pressures. **B**, Plot of *v*/[E_0_](s^−1^) over entire pressure and temperature range of HRP, data was recorded in triplicate. *Conditions*, 0.2 *n*M HRP, 1.8 mM H_2_O_2_ and 73 mM ABTS in 50 mM HEPES buffer, pH 6.5.

